# Bioorthogonal Tuning of Hydrogel Stiffness Promotes Zonal Redifferentiation of Passaged Chondrocytes

**DOI:** 10.64898/2026.07.02.736090

**Authors:** Thomas J. Manzoni, Aditya Natu, John E. Caputo, Anh Ho, Isabella Ewine, Lilly Smull, Yinzhi Fang, Joseph M. Fox, Alvin W. Su, Xinqiao Jia, Justin Parreno

**Author notes:** Denotes Co-Corresponding Author, Correspondence to: Justin Parreno, Wolf Hall, 105 The Green, Newark, Delaware, 19716, USA, Xinqiao Jia, 201 DuPont Hall, 127 The Green, Newark, Delaware, 19716, USA. Denotes Co-First Author (Thomas J. Manzoni, Aditya Natu).

## Abstract

Generating bioengineered cartilage that recapitulates the depth-dependent phenotype, structure, and function of native articular cartilage remains a challenge. While cartilage is rich in aggrecan and type II collagen, proper function depends on depth-dependent protein expression. Superficial zone chondrocytes (SZCs) secrete proteoglycan-4 (PRG4) to lubricate the cartilage surface. Deep zone chondrocytes produce type X collagen (COLX) to support compressive loading and load transfer to subchondral bone. We previously demonstrated that passaged full-thickness chondrocytes (FTCs) and zonal chondrocytes can re-express cartilage and zone-specific markers following scaffold-free three-dimensional (3D) culture in redifferentiation media. However, in the absence of an instructive matrix, cells expressed low levels of zone-specific proteins and exhibited limited depth-dependent organization. *We hypothesize that synthetic extracellular matrix with zone-specific microenvironmental cues will guide zonal differentiation*. To this end, passaged primary bovine chondrocytes were encapsulated in a soft, hyaluronan (HA)-based, cell-adhesive, and protease-degradable hydrogel established via bioorthogonal tetrazine (Tz) ligation with norbornene (Nb). When supplemented with TGFβ3, FTCs deposited aggrecan and type II collagen with minimal type I collagen. Application of interfacial tetrazine ligation with *trans*-cyclooctene (TCO) during cell culture resulted in matrix stiffening, leading to upregulation of COLX expression. Conversely, SZCs cultured in soft hydrogels exhibited the greatest PRG4 expression. Establishment of a trilayered construct with region-specific stiffness via the diffusion-controlled reaction promoted PRG4 and COLX expression in defined zones. Together, these findings demonstrate that tunable HA-based hydrogels can enhance zone-specific chondrocyte phenotypes and promote the formation of zonally organized cartilage.

## INTRODUCTION

Articular cartilage is a highly specialized connective tissue that facilitates the transmission and distribution of mechanical load with a low coefficient of friction during joint articulation. Maintained by chondrocytes, the articular cartilage matrix is comprised primarily of proteoglycans (Aggrecan; ACAN), collagens (Collagen II; COL2), hyaluronan (HA), and water. The extracellular matrix is spatially organized into three distinct zones, the superficial zone (SZ), middle zone (MZ), and deep zone (DZ), each with unique characteristics [1]. Owing to its avascular nature, when damaged, articular cartilage is incapable of self-repair [1, 2]. Cell-based strategies have been developed to repair small cartilage defects by directly implanting passaged chondrocytes into the damaged region for localized tissue repair [3–5]; the goal is to regenerate an ECM that mimics the composition and organizational properties of native articular cartilage.

Successful regeneration of zonal cartilage requires the restoration of zone-specific chondrocyte phenotypes, as cells within each zone are responsible for producing and maintaining the specialized extracellular matrix that contributes to the proper function of the tissue. Superficial zone chondrocytes (SZCs) are responsible for producing lubricin/proteoglycan-4 (PRG4), which lubricates the articular surface, establishing a low coefficient of friction [6–8]. Chondrocytes in MZ produce cartilage intermediate layer protein (CILP), a matrix-associated protein that contributes to extracellular matrix structural integrity [9]. Deep zone chondrocytes (DZCs) express alkaline phosphatase (ALP), chondroadherin (CHAD), and collagen type X (COLX), all of which contribute to matrix mineralization and mechanical integration with the subchondral bone [10–14]. Furthermore, glycosaminoglycan (GAG) content increases from SZ to DZ, leading to an intrinsic gradient in tissue stiffness [15, 16]. Collectively, these zone-specific matrix properties are essential for maintaining tissue function.

Passaged chondrocytes are a common FDA-approved cell source for cell-based cartilage repair. To obtain a sufficient number of cells, primary chondrocytes are expanded on tissue culture polystyrene as a two-dimensional (2D) monolayer. During 2D expansion, chondrocytes undergo dedifferentiation, characterized by reorganization of cortical F-actin into stress fibers [17, 18]. These morphological changes coincide with a loss of expression of cartilage matrix markers and an increase in expression of fibroblastic matrix components, including type I collagen (COL1) and tenascin C (TNC), contractile proteins such as α–smooth muscle actin (αSMA) [17, 19–22]. Consequently, the dedifferentiated chondrocytes form mechanically inferior cartilage [23, 24]. It has been shown that full-thickness chondrocytes (FTCs) can be redifferentiated to form bioengineered cartilage high in ACAN and COL2 content [25–27]. However, although redifferentiated FTCs support the formation of bulk cartilage matrix, they do not fully restore the zonal characteristics of native articular cartilage, particularly with regard to the expression of SZ marker PRG4. To address this limitation, we developed strategies to enhance the expansion and redifferentiation of SZC, improving their ability to re-establish zone-specific phenotypes following passaging [28, 29]. Despite these advances, current approaches are limited in their ability to recapitulate the zonal organization and depth-dependent expression of native articular cartilage.

Three-dimensional culture of chondrocytes in customized hydrogel matrices with defined extracellular cues offers a promising strategy to address this limitation. Hydrogel stiffness has been shown to play a critical role in chondrocyte phenotype and matrix deposition [30]. However, reported hydrogel systems fail to mimic the dynamic mechanical microenvironment of the developing joint. During early cartilage development, a primitive, hyaluronan (HA)-rich matrix undergoes rapid stiffening that precedes and drives both matrix deposition and spatial organization [31], resulting in mature cartilage with depth-dependent stiffness [32, 33]. This suggests that dynamic matrix stiffening may influence zonal chondrocyte phenotype; however, whether biomechanical cues can promote zonal redifferentiation in a bioengineering context remains unknown.

We previously reported a bioorthogonally tunable hydrogel derived from HA, hereafter referred to as BOHAgel, decorated with cell-adhesive RGD motifs, and crosslinked with a matrix metalloproteinase (MMP)-degradable peptide through tetrazine (Tz) ligation with norbornene (Nb). Spatiotemporal control over matrix properties, including stiffness and adhesiveness, is achieved during cell culture through a cycloaddition reaction between the tetrazine moieties and the *trans*-cyclooctene (TCO) species at the gel-liquid interface [34–37]. With the post-gelation modification being diffusion-controlled, region-specific microenvironments that more closely resemble native cartilage zones can be established by controlling the diffusion of the TCO species [35]. Importantly, this capability aligns with the rapid stiffening of a primitive, HA-rich matrix observed during early cartilage development [36]. Therefore, BOHAgel provides a promising platform to recapitulate these dynamic mechanical cues to drive chondrocyte redifferentiation. This study aims to determine whether BOHAgel can serve as an artificial matrix for generating zone-specific microenvironmental cues. *We hypothesize that 3D culture of FTCs or SZCs in dynamically tunable BOHAgel will yield bioengineered cartilage with zonal expression*. This study may provide a strategy to better recapitulate native-like bioengineered tissue, thereby improving cartilage repair outcomes.

## METHODS

### Isolation and Culture of FTCs

Full-thickness chondrocytes were isolated from bovine metacarpal-phalangeal joints as previously described [18, 25]. Briefly, the full depth of cartilage tissue up to the subchondral bone was manually dissected from the joint. Chondrocytes were isolated by serial enzymatic digestion in 0.5% Protease XIV from *Streptomyces griseus* (Sigma-Aldrich, St. Louis, MO, USA) for 45 min at 37°C, followed by 0.1% Collagenase A from *Clostridium histolyticum* (Roche, Mannheim, Germany) for 14–17 h at 37°C. For monolayer expansion, FTCs were seeded at 1,500 cells/cm² on tissue culture polystyrene (GenClone, Genesee Scientific, San Diego, CA, USA). Cells were expanded in Ham’s F-12 medium (Corning, Edison, New Jersey, USA) supplemented with 10% fetal bovine serum (GenClone) and 1% antibiotic-antimycotic (Corning). Cells were cultured to approximately 80–90% confluence and then detached with 0.25% trypsin-EDTA (GenClone), generating passage 1 (P1) cells. P1 FTCs were reseeded at 2×10³ cells/cm² on polystyrene. P1 cells were cultured to 80–90% confluence and detached to obtain passage 2 (P2) cells.

### Isolation and Enrichment of SZCs

SZ cartilage was manually microdissected from the joint using a surgical scalpel as previously described [28, 29, 38]. SZC were isolated by serial enzymatic digestion in 0.5% Protease XIV from *Streptomyces griseus* for 45 min at 37°C, followed by 0.1% Collagenase A from *Clostridium histolyticum* for 14–17 h at 37°C. For monolayer expansion, SZC were seeded at 1,500 cells/cm² on polystyrene coated with cartilage-derived extracellular matrix (CD-ECM; StemBioSys, San Antonio, TX, USA) and maintained in Ham’s F-12 medium supplemented with 10% fetal bovine serum and 1% antibiotic-antimycotic. At approximately 80–90% confluence, cells were detached with 0.25% trypsin-EDTA, generating P1 cells. P1 SZC were reseeded at 2×10³ cells/cm² on CD-ECM. Cells were again cultured to 80–90% confluence and detached to obtain P2 cells.

### Chondrocyte Encapsulation and 3D Culture

Hydrogel precursors, including tetrazine-functionalized HA (HA-Tz), Nb-tagged MMP-degradable crosslinker (SMR-bisNb), and TCO-conjugated GRGDSP (RGD-TCO), were synthesized following our previously reported procedures.[35]. TCO-modified HA (HA-TCO) was prepared by treating 5 kDa HA with a hydrazide-modified TCO in H_2_O/DMSO (see Supporting Information). Stock solutions of HA-Tz and RGD-TCO in calcium- and magnesium-free PBS (PBS-/-; Quality Biological, Gaithersburg, MD, USA) were mixed and vortexed before P2 FTCs or SZCs were introduced and homogeneously suspended. Cellular constructs were produced upon addition of SMR-bisNb in PBS at a Tz/Nb molar ratio of 5/2. The hydrogel mixture (75 or 200 μL) was aliquoted on a 0.4-µm cell culture insert (Cell Treat, Ayer, MA, USA) and incubated at 37°C for 45 min before the addition of redifferentiation media (2 mL) consisting of Dulbecco’s modified Eagle medium (GenClone) supplemented with ITS+ Premix (Corning) containing 6.25 μg/mL insulin, 6.25 μg/mL transferrin, 6.25 μg/mL selenium, 1.25 mg/mL BSA, and 5.35 ng/mL linoleic acid, 0.1 μM dexamethasone (Sigma Aldrich), 40 μg/mL proline (Sigma Aldrich), 110 μg/mL pyruvate, and 100 μg/mL ascorbic acid (Sigma Aldrich). In some cases, the redifferentiation media was supplemented with 10 ng/mL transforming growth factor beta 3 (TGFβ3; R&D Systems, Minneapolis, MN, USA) on day 7 of culture. Monolithic constructs were prepared with 0.5 mM RGD and 10 or 20 mg/mL HA-Tz. To prepare a multilayered construct, a hydrogel mixture (60 μL) containing 20 mg/mL HA-Tz and 0.5 mM RGD was overlaid on a partially cross-linked gel (30 µL) containing 10 mg/mL HA-Tz and 0.5 mM RGD. Thirty minutes later, media was added to the well, and 3D cultures were maintained for 7 days. On day 7, media was supplemented with HA-TCO (20 mg/mL), and interfacial bioorthogonal crosslinking was allowed to proceed for 24 h or 45 min to yield homogeneously stiffened or trilayered constructs, respectively. Subsequently, chondrocyte cultures were maintained in HA-TCO-free media, with media refreshment every two days.

### Oscillatory Rheology

Fully swollen acellular hydrogels (200 *µ*L, 10 or 20 mg/mL HA) with or without interfacial stiffening were loaded onto an 8-mm parallel plate geometry on a Discovery Hybrid Rheometer (DHR3, TA Instruments, Newcastle, DE). Mineral oil was applied around the geometry to prevent sample dehydration during measurements. After application of an axial force of 0.015 N, a frequency sweep was carried out from 0.05 to 10 Hz at 5% strain at 25°C. The storage (G’) and loss (G’’) moduli are reported as average values from three replicates.

### Confocal Analysis of Tri-layered Hydrogel

To enable fluorescent labeling, 10 and 20 mg/mL HA-Tz were mixed with Alexa Fluor 488-TCO and Alexa Fluor 568-TCO (5 µM; Click Chemistry Tools), respectively. Separately, HA-TCO (20 mg/mL) was reacted with Cy5-tetrazine (5 µM) by vortex mixing. Hydrogels were prepared using fluorescently labeled precursors following the procedure described without cells. Interfacial Tz-TCO ligation was conducted for 0, 15, 30, 45, or 60 min at 37°C. At each point, the HA-TCO solution was aspirated, and the construct was washed three times with PBS to terminate the diffusion-controlled stiffening process prior to imaging. Confocal imaging was performed using a Zeiss LSM 880 laser scanning confocal microscope with a 5× objective (NA: 0.25). Images were taken using a Z-stack spanning 1.5 mm at 20 μm step intervals. FIJI was used for line-scan analysis. Images were imported and split into individual channels corresponding to Alexa Fluor 488, Alexa Fluor 568, and Cy5 signals. A line with a fixed width of 500 pixels was drawn perpendicular to the hydrogel interface and applied across all samples. Fluorescence intensity for each channel was measured along the line scan and plotted as a percentage distance across the hydrogel. To quantify the percentage of the hydrogel that was stiffened, the Cy5 fluorescence peak was identified from the line scan, and the distance from the hydrogel surface to the Cy5 peak was divided by the total hydrogel thickness.

### Live/Dead Imaging and Quantification

Cellular constructs were washed with PBS and incubated in the dark in a staining solution (2 mL) consisting of Hoechst 33342 (1:500; 1mg/mL; Biotium, Fremont, CA, USA), Calcein-AM (1:1000; Biotium), and Ethidium Homodimer-1 (EthD-1:1000; Biotium) at 37°C for 30 min. After three PBS washes, the construct was cut in half and placed face down on a Zeiss LSM 880 laser-scanning confocal microscope (Zeiss, Jena, Germany) with a 20× objective (NA: 0.4; 1-μm slices). Multiple fields of view were imaged per sample, and Z-stacks were produced. Using FIJI (ImageJ; National Institutes of Health), percent viability was calculated as the difference between Hoechst-positive nuclei and EthD-1–positive nuclei, relative to all Hoechst-positive cells.

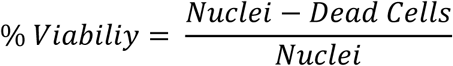

### F-Actin Visualization and Quantification

After overnight fixation in 4% paraformaldehyde (Electron Microscopy Sciences, Hatfield, PA, USA) at 4°C, constructs were washed with PBS and cryoprotected in 30% sucrose (Sigma-Aldrich) overnight at 4°C. Samples were then embedded in OCT compound (Sakura Finetek, Torrance, CA, USA), snap-frozen, and cryosectioned at a thickness of 20 μm. After blocking with 20% goat serum, sections were incubated with PBS containing Hoechst 33342 and rhodamine-phalloidin (1:50; Biotium) for 24 h at 4°C. Samples were then washed with PBS three times and mounted with rectangular No. 1.5 glass coverslips (Globe Scientific Inc., Mahwah, NJ, USA) using Drop-n-Stain mounting medium (Biotium). Fluorescence imaging was performed using a Zeiss LSM 880 laser-scanning confocal microscope. Z-stack images were acquired using either 40× (NA: 1.3 oil immersion; 0.4 μm step size) or 63× (NA: 1.4 oil immersion; 0.3 μm step size) objectives. Images were processed using Zen Blue (Zeiss). Using 40× images, encapsulated cells were manually traced based on rhodamine-phalloidin staining using FIJI. For each tissue section, 20 cells were analyzed per field, and this was repeated across three sections from different tissues. Cell area and circularity were quantified as previously described [18, 39]. In total, a minimum of 60 cells were analyzed per condition.

### RNA Extraction and RT-PCR

Total RNA was isolated using TRIzol reagent (Sigma-Aldrich) in accordance with the manufacturer’s instructions. RNA concentration and purity were determined using a NanoDrop One spectrophotometer (Thermo Fisher Scientific, Waltham, MA, USA) based on A260/A280 and A260/A230 absorbance ratios. Samples were reverse transcribed to complementary DNA (cDNA) using UltraScript 2.0 cDNA Synthesis Kit (PCR Biosystems; Wayne, PA, USA) according to the manufacturer’s directions. Quantitative real-time PCR (qPCR) was performed using 20 ng of cDNA per reaction with gene-specific primers (10 μM working concentration) and qPCRBio SyGreen Blue Mix (PCR Biosystems). The primer information can be found in Table S1. All primer pairs were designed using NCBI Primer-BLAST and were previously validated [19, 40]. Reactions were run in technical duplicate on a Cielo 3 Real-Time PCR System (Azure Biosystems, Houston, TX, USA). Relative gene expression was calculated using the Pfaffl method [41], with 18S rRNA serving as the endogenous housekeeping gene. Expression levels were reported as a percentage relative to the 2D monolayer controls.

### Immunostaining, Imaging, and Quantification

Cellular constructs were fixed and cryosectioned at a thickness of 20 μm. For antigen retrieval, sections were digested in 0.4% pepsin (Sigma-Aldrich) in 1× Tris-buffered saline (pH 2) for 5 min, then blocked with 20% goat serum. Sections were then incubated overnight at 4°C with primary antibodies (Table S2A). The next day, sections were washed three times with PBS, incubated with secondary antibodies (Table S2B), and counterstained with Hoechst 33342 for 1 h at room temperature, with three PBS washes between each step. Slides were prepared using Drop-n-Stain mounting and glass coverslips. Confocal imaging was performed using a Zeiss LSM 880 laser scanning confocal fluorescence microscope with a 5× objective (NA: 0.25; 1-μm slices). Z-stack images were acquired with a step size of 1 μm. Images were processed using Zen Blue (Zeiss), and the mean fluorescent intensity (MFI) was determined using FIJI. Tissue boundaries were manually traced, and the MFI of the channel corresponding to the protein of interest was measured. MFI values were normalized to the average native Bovine cartilage control within each experimental set and subsequently pooled for analysis. For quantification of the expression levels of zone-specific markers, MFI values were compared to region-matched native tissue controls, defined as the top 20% of native cartilage for SZ proteins, the middle 40% for MZ proteins, and the bottom 40% for DZ proteins. To analyze the trilayered constructs, tissue sections were divided along the depth of the hydrogel into three regions based on relative thickness: the top 10% (10 mg/mL HA region), the middle 45% (20 mg/mL region), and the bottom 45% (stiffened 20 mg/mL HA region). The MFI of the channel corresponding to the protein of interest was measured within each ROI. MFI values were normalized to the top 10% region within each sample and expressed as relative values.

### Metabolic Labeling of Newly Synthesized GAGs and Collagens

To label newly synthesized collagens via copper-free click chemistry [42], 48 h before the culture was terminated, the standard media was replaced with modified depletion media consisting of DMEM-LM (ThermoFisher) supplemented with 105 μg/mL L-leucine (Sigma-Aldrich), standard culture supplements, TGF-β3 and 30 μM L-azidohomoalanine (AHA; Vector Laboratories, Newark, CA, USA). Next, samples were incubated with 647-DBCO (Vector Laboratories) in PBS containing 1% bovine serum albumin (Prometheus Biosciences, San Diego, CA, USA) for 1 h before imaging. Similarly, 24 h before the culture was terminated, the media was supplemented with 30 μM N-azidoacetylgalactosamine-tetraacylated (GAL; Vector Laboratories). Samples were then incubated with 647-DBCO in PBS containing 1% bovine serum albumin for 1 h to fluorescently label newly synthesized GAG. For visualization, labeled constructs were counterstained with Calcein-AM (1:1000) and imaged with a Zeiss LSM 880 laser-scanning confocal microscope. Z-stack images were acquired using a 40×/1.3 NA oil objective with a 0.7 μm step size. MFI was quantified as described above. For quantification, samples were digested in PBS containing 0.1% collagenase and 100 U/mL hyaluronidase. The fluorescence intensity of the digested samples was measured using a Promega GloMax Discover plate reader (Excitation 627 nm, Emission 660-720 nm; Promega, Madison, WI, USA). Fluorescence values were normalized to the wet weight of each sample and expressed as a percentage of the corresponding control.

### Histology

Samples harvested on day 21 were fixed in 10% neutral buffered formalin (StatLab, McKinney, TX, USA; pH 7.4) overnight at 4°C. Samples were then dehydrated through a graded ethanol series, cleared in xylene, embedded in paraffin, and sectioned at 5 μm thickness. Sections were stained with hematoxylin and eosin (H&E) to assess morphology, toluidine blue to evaluate GAG content, or picrosirius red to visualize collagens before mounting the coverslip. Brightfield images were acquired using a Zeiss Axio Observer (Zeiss) with a 2.5× objective (NA: 0.12).

### Statistical Analysis

All experiments were repeated at least three times, with each independent experiment performed using cells pooled from multiple bovine legs. Data from individual experiments were aggregated for analysis. Statistical analyses were conducted using GraphPad Prism (GraphPad Software Inc., Boston, MA, USA). Outliers were identified using the ROUT method [43]. For comparisons between two groups, an unpaired t-test was used. For comparisons involving more than two groups, a one-way analysis of variance (ANOVA) was performed to assess differences among group means, followed by planned comparisons using the Bartlett correction method to identify differences between groups. Data presented as mean ± standard deviation.

## RESULTS

### Rapid Tz-TCO cycloaddition reaction enables bioinspired spatiotemporal stiffening of BOHAgels

Monolithic BOHAgels containing unreacted tetrazines were prepared via the Tz/Nb reaction at a molar ratio of 5/2 (Figure 1A). Rheological testing of gels made with 10 mg/mL (BOHAgel-1) or 20 mg/mL (BOHAgel-2) HA-Tz revealed an average shear elastic modulus (G’) of 108 Pa and 437 Pa, respectively (Figure 1B). Consumption of the remaining Tz groups in the networks via interfacial crosslinking using HA-TCO (Figure 1A) resulted in an increase in G’ (174 Pa for BOHAgel-1 and 975 Pa for BOHAgel-2, Figure 1B, Figure S2). In all cases, the loss tangent, i.e., G”/G’, was in the range of 0.01-0.03, indicative of a highly elastic network dominated by covalent crosslinks [35].

**Figure 1:**
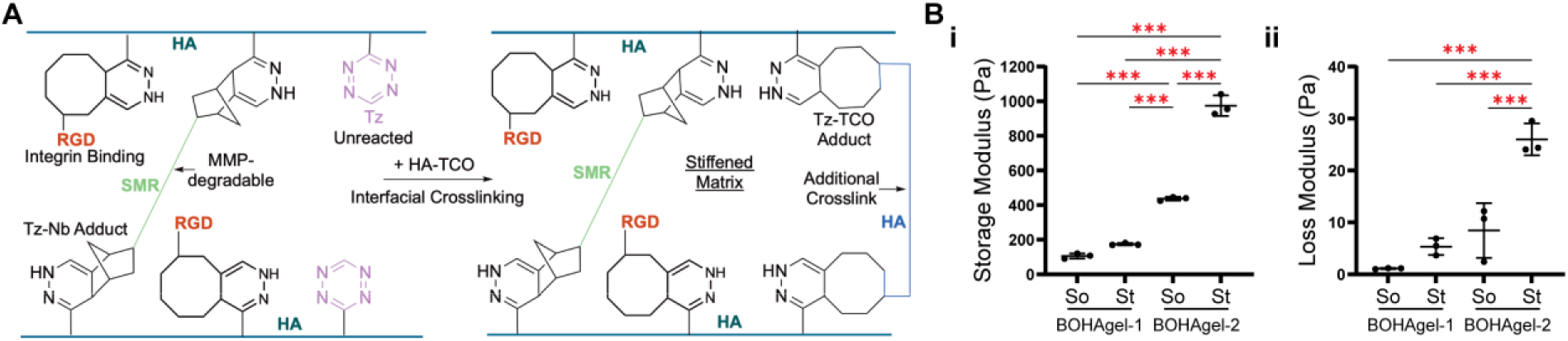
BOHAgels can be dynamically tuned during cell culture to enable in situ matrix stiffening. (**A**) Reaction scheme showing the parent BOHAgel established via Tz/Nb cycloaddition with Tz in stoichiometric excess. Post-gelation modification was achieved via the Tz/TCO cycloaddition which is faster than molecular diffusion. (**B**) Storage (i) and loss (ii) moduli of BOHAgel-1 (containing 1 wt% HA-Tz) and −2 (containing 2 wt% HA-Tz) with (stiffened: St) or without (soft: So) interfacial crosslinking. ***p < 0.001.

To generate tri-layered scaffolds, we initially layered BOHAgel-2 on top of BOHAgel-1. We then allowed the diffusion-mediated crosslinking to occur through the top 50% of the gel volume. To visualize the evolution of matrix composition and mechanics, we fluorescently labeled the HA-Tz component in BOHAgel-1 and BOHAgel-2 with Alexa Fluor-448 and Alexa Fluor 568, respectively, and HA-TCO with Cy5 (Figure 2A). Using confocal imaging with line-scan analysis, we revealed a gradual increase in the depth of the stiffened region incorporating HA-TCO over time (Figure 2B–D). When the interfacial reaction was allowed to proceed for 45 min, 40%-50% of the scaffold was stiffened. Note that the fluorescent signals from the lower half of the scaffold remain unchanged, indicating that 45 min is not sufficient for HA-TCO to diffuse into these regions. In addition, the percentage of hydrogel stiffening remains stable over 14 days of incubation (Figure S3). Collectively, tetrazine ligation enables the development of a multilayered scaffold (Figure 2E) with region-specific mechanical cues.

**Figure 2:**
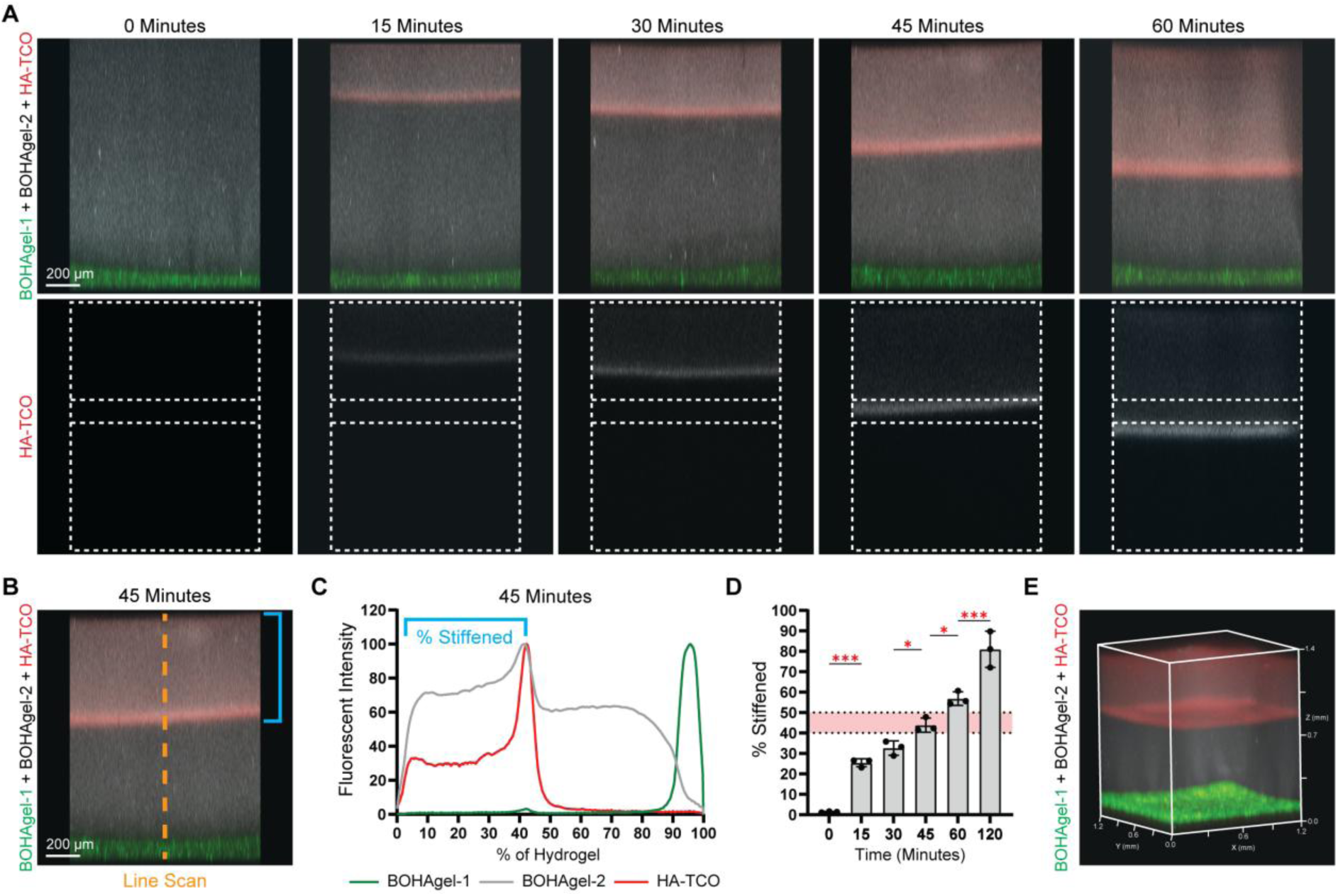
Application of diffusion-controlled interfacial crosslinking resulted in the establishment of trilayered construct with depth-dependent stiffness. **(A)** Confocal microscopy images showing time-dependent incorporation of HA-TCO (red) in the top BOHAgel-2 (gray). BOHAgel-1 (green) remained intact. **(B-D)** Representative line scan analyses **(B-C)** and quantification **(D)** of the percentage of the hydrogel that had been stiffened. **(E)** 3D reconstructed confocal image illustrating the formation of a trilayered scaffold following 45 min incubation with HA-TCO. *p < 0.05, ***p < 0.001.

### BOHAgels prime passaged chondrocytes for chondrogenic redifferentiation

Passaged FTCs were encapsulated in BOHAgel-1 and maintained in differentiation media for up to 42 days. The live/dead assay (Figure 3A) showed that encapsulated cells were 99.2% viable on day 0. Although compared to day 0, there was a moderate decrease in cell viability on days 7, 21, and 42, >88% of cells remained viable throughout 42 days of culture. F-actin staining revealed that, compared to day 0, cells on day 7 were larger (Figure 3B). Cell area decreased with time in culture (day 7 and 14) and became similar in size to cells at day 0. Cell circularity was higher on day 0 than on days 7, 14, and 21. Throughout culture, cells exhibited a cortical F-actin organization consistent with a rounded morphology (Figure 3C). By day 21, while cells remain round, they contain F-actin protrusions.

**Figure 3:**
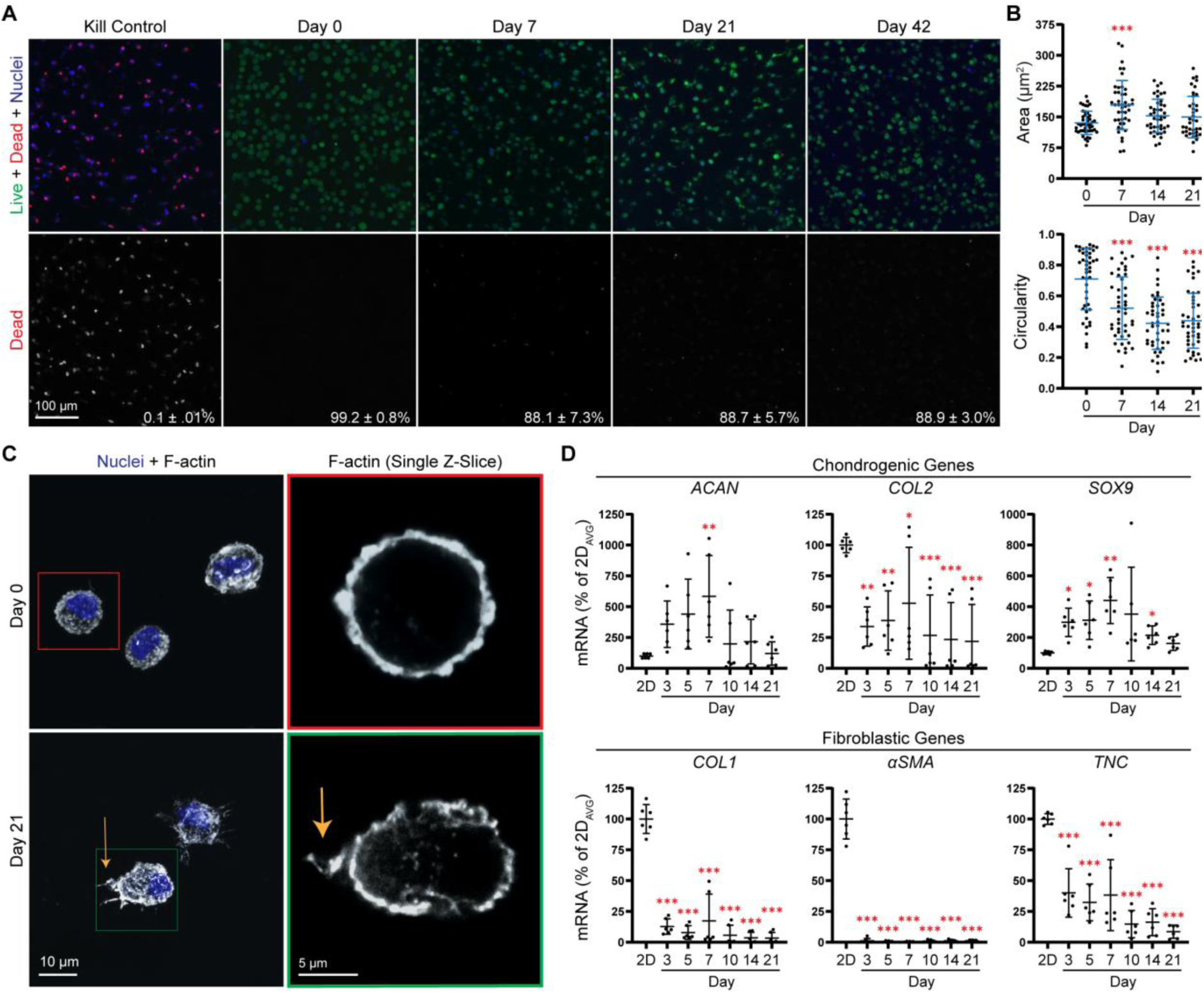
BOHAgel-1 primes passaged FTCs cells for redifferentiation. **(A)** Confocal microscopy images of cellular constructs after 0, 7, 21, and 42 days of culture. Green: Calcein-AM for live cells; Red (Top Panel) and Gray (Bottom Pannel): Ethidium Homodimer-1 for dead cells; Blue: Hoechst for Nuclei. The corresponding viability is indicated in the bottom-right corner of each image. **(B)** Quantification of cell area and circularity. **(C)** Confocal microscopy images of chondrocytes stained for F-actin (Grey) and nuclei (blue) at day 0 and 21 of culture. Orange arrows indicate F-actin protrusions. **(D)** Real-time RT-PCR analysis of mRNA levels of chondrogenic and fibroblastic markers expressed by encapsulated chondrocytes. Expression was normalized to the 2D control as a percentage. *p < 0.05, **p < 0.01, ***p < 0.001.

Next, we evaluated mRNA levels of chondrogenic and fibroblastic matrix markers over 21 days (Figure 3D). For chondrogenic markers, *ACAN* expression increased significantly on day 7, but decreased by day 14. Compared with P2 cells in 2D culture, cells maintained in 3D expressed lower levels of *COL2* at all time points. The mRNA levels for SRY-Box Transcription Factor 9 (*SOX9*), a transcriptional activator of *ACAN* and *COL2* [44, 45], increased at days 3, 5, 7, and 10 relative to the 2D P2 cells. Conversely, culture in BOHAgels decreased fibroblastic *COL1, αSMA,* and *TNC* mRNA levels throughout culture. Collectively, BOHAgels culture of passaged FTCs led to an increased expression of chondrogenic markers while repressing the fibroblastic phenotype.

### Soluble TGFβ3 supplemented to 3D cultures further enhances chondrogenic redifferentiation

While our data show that BOHAgel-1 primes P2 FTCs for redifferentiation, additional soluble factors may be required to increase *COL2* mRNA levels, thereby strengthening lineage commitment. Since TGFβ3 supplementation has previously been shown to promote cartilage matrix deposition by passaged cells [25], we next tested its applicability in our 3D cultures. Here, TGFβ3 was supplemented in the media from day 7 of culture. Our results (Figure 4A-C) show that TGFβ3 treatment did not significantly alter cell morphology, F-actin organization, cell area, or circularity. Under both conditions, cells remained round, displaying cortical F-actin and exhibited protrusions. On the other hand, there was a significant increase in *ACAN* and *COL2* mRNA levels at day 14 in TGFβ3-treated groups relative to both 2D and 3D untreated controls (Figure 4D). By day 21, *ACAN* and *COL2* mRNA levels in TGFβ3-treated cells decreased to the levels observed in untreated controls. There were no differences in *SOX9* mRNA levels. While TGFβ3 does not alter *TNC* mRNA levels, TGFβ3 slightly increases *COL1* and *aSMA* mRNA levels on day 14. However, by day 21 there are no differences in *COL1*, *aSMA,* and *TNC* mRNA levels between TGFβ3 treated and untreated cells. Additionally, *COL1*, *αSMA*, and *TNC* mRNA levels remain significantly lower in both treated and untreated groups relative to 2D controls. Thus, TGFβ3 enhanced the mRNA levels of chondrogenic genes in passaged FTCs maintained in BOHAgels.

**Figure 4:**
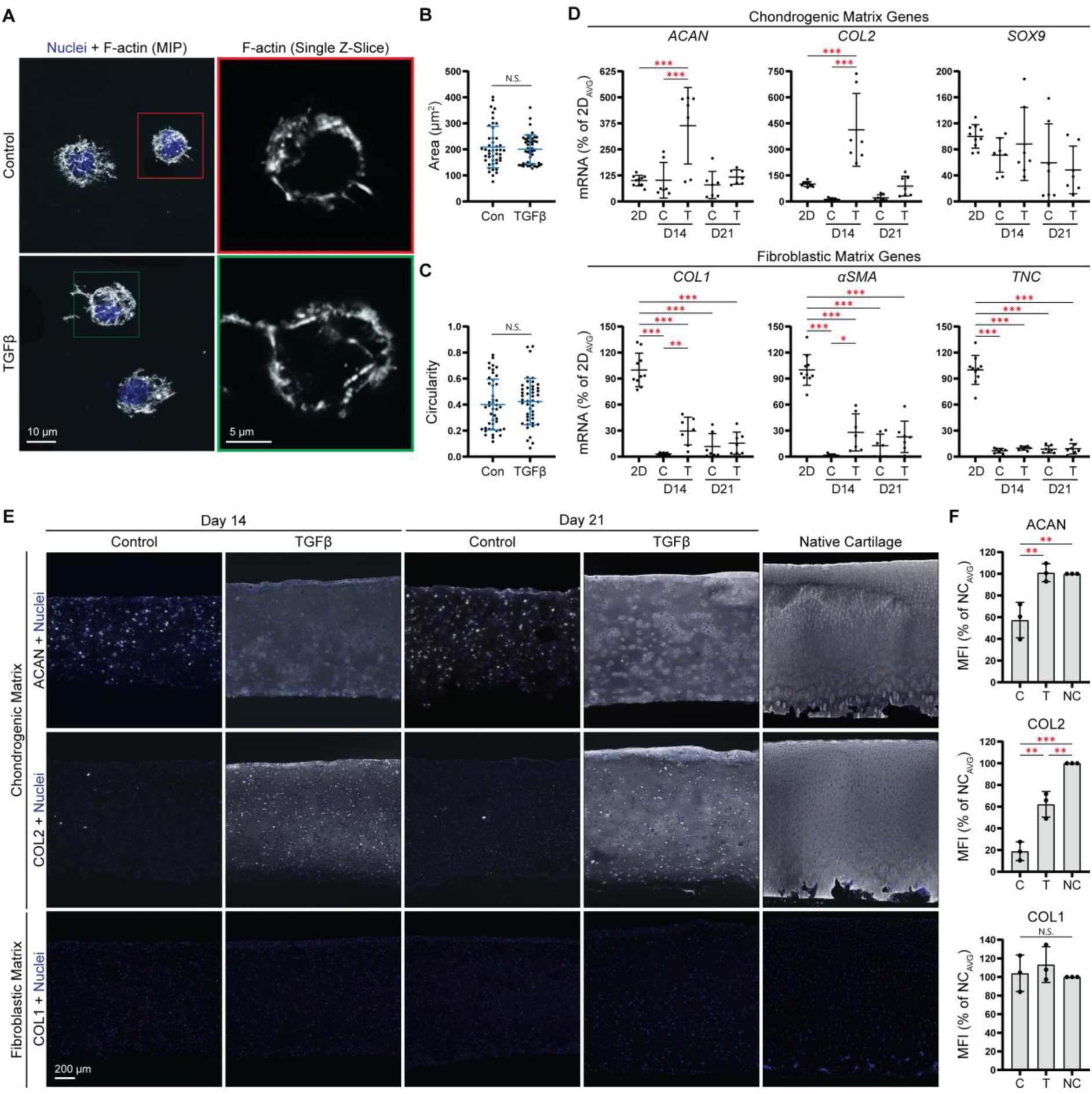
Passaged chondrocytes maintained in BOHAgel-1 produce cartilage-like matrix when treated with TGFβ3. **(A)** Confocal microscopy images showing F-actin (grey) organization. **(B-C)** Quantification of cell area **(B)** and circularity **(C)** at day 21 of culture. **(D)** Real-time RT-PCR analysis of mRNA levels of chondrogenic and fibroblastic markers expressed by resident FTCs in control media (C) or TGFβ3-supplemented (T) media. The expression is normalized to 2D controls, as a percentage. **(E)** Confocal microscopy images 3D cultures showing the accumulation of chondrogenic proteins ACAN and COL2 and fibroblastic matrix protein COL1. Specific proteins are shown in grayscale and counterstained with Hoechst for Nuclei (Blue). **(F)** Quantification of protein production by mean fluorescent intensity (MFI). N.S.: not Significant), *p < 0.05, **p < 0.01, ***p < 0.001.

At the protein level by immunofluorescence (Figure 4E), TGFβ3 treatment increased the accumulation of cartilage-specific matrix proteins ACAN and COL2 at both day 14 and day 21, compared to untreated controls. Notably, the MFI for ACAN in TGFβ3-treated constructs is similar to that for the native articular cartilage (Figure 4F), whereas the untreated controls was significantly lower. Although TGFβ3 enhanced the staining intensity of COL2, COL2 content remained lower than that of native cartilage. In contrast, COL1 intensity remained low across all conditions and did not differ significantly between groups. Collectively, these findings demonstrate that TGFβ3 promotes cartilage-like matrix deposition by passaged FTCs maintained in BOHAgels. Therefore, TGFβ3 was used to stimulate redifferentiation and cartilage matrix deposition in subsequent experiments.

### Matrix stiffening promotes cortical actin and increases chondrogenic redifferentiation

We sought to evaluate the effects of dynamic stiffening on chondrocyte phenotype and matrix production. To this end, BOHAgel-2 was used to maximize the magnitude of the stiffness change following crosslinking. Specifically, P2 FTCs were encapsulated in BOHAgel-2 with an initial G’ of 438 Pa. After 7 days of culture, HA-TCO and TGFβ3 were both supplemented in the media. Twenty-four hours later, excess HA-TCO was depleted, and cells were maintained in a stiffened environment (G’ 978 Pa) in TGFβ3 media for an additional 14 days. F-actin staining shows that matrix stiffening maintained the cortical F-actin organization, however, blunted the F-actin protrusions previously observed in unstiffened BOHAgel-1 (Figure 5A). In addition, stiffening did not affect cell circularity but reduced cell area (Figure 5B, C). At the mRNA level (Figure 5D), matrix stiffening increased *ACAN* levels relative to both 2D controls and the soft BOHAgel-2 counterpart on day 21; a similar increase in *COL2* transcript was detected on both days 14 and 21. Matrix stiffening did not alter the expression of *SOX9* at either time point, nor did it affect the expression of genes encoding *COL1*, *αSMA*, and *TNC*. Notably, compared with cells in 2D monolayer culture, the mRNA levels of COL1, αSMA, and TNC remained significantly lower in P2 FTCs maintained in Orthgel20, regardless of whether the matrix had been stiffened.

**Figure 5:**
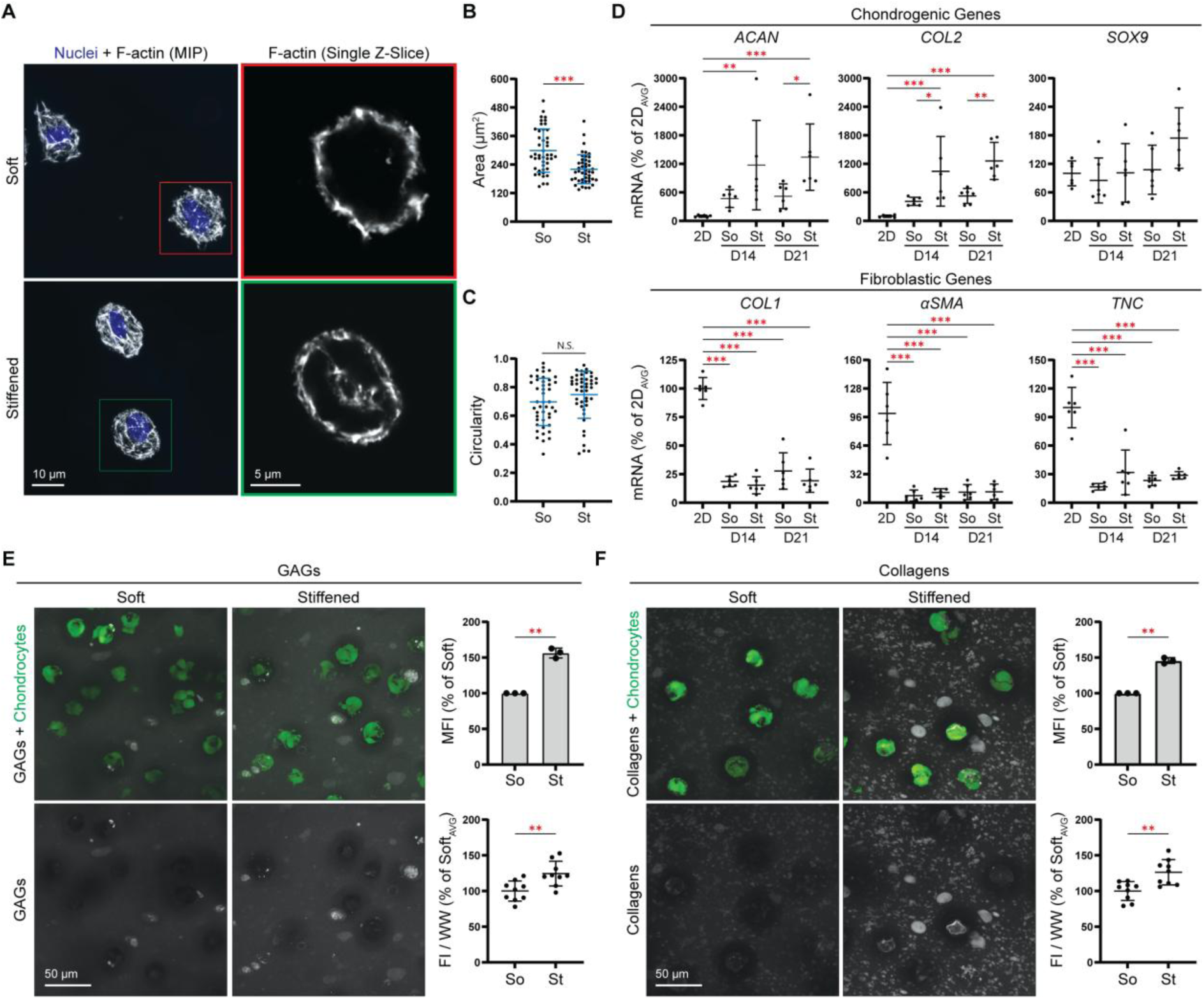
Matrix stiffening promotes cortical actin organization and enhances matrix deposition by FTCs in BOHAgel-2. **(A)** Confocal microscopy images of cells maintained in the parent soft (So) and the stiffened (St) hydrogels after phalloidin staining. (**B-C**) Quantification of cell area **(B)** and (C) circularity for day 21 cultures. **(D)** Real-time RT-PCR analysis of mRNA levels of chondrogenic and fibroblastic markers. Data were normalized to 2D controls and expressed as percentages. (**E-F**) Characterization of GAGs accumulated on days 20-21 **(E)** and collagens accumulated on days 19-21 **(F)** through Click-chemistry based metabolic labeling. Labeled proteins are shown in gray, and chondrocytes are stained with Calcein (green). N.S.; not significant, *p < 0.05, **p < 0.01, ***p < 0.001.

Next, we employed Click chemistry-based metabolic labeling to quantify newly synthesized GAGs and collagens produced by cells residing in BOHAgel-2 (Figure 5E–F). Here, GAG labels were metabolically incorporated in cells between days 20–21 of culture, whereas collagen labels were incorporated between days 19–21. Compared with those maintained in the soft counterparts, cells cultured in the stiffened environment produced higher levels of GAGs and collagens. These results demonstrate that matrix stiffening enhances chondrogenic differentiation of P2 FTCs within dynamically tunable 3D BOHAgels.

### Matrix stiffening increases DZ-associated expression by passaged FTCs

We then inquired whether dynamic matrix stiffening stimulates FTCs to express MZ and DZ-associated matrix molecules. At the transcript level, compared with cells cultured in 2D, FTCs maintained in BOHAgel-2 without interfacial crosslinking showed higher levels of the MZ marker *CLIP*; matrix stiffening led to further increase (Figure 6A). Culture in unmodified BOHAgel-2 increased *CHAD* and *ALP* expression relative to 2D controls but did not alter *COLX* expression. Day 7 matrix stiffening increased *COLX*, *CHAD*, and *ALP* expression relative to 2D controls. Compared with cells residing in unmodified BOHAgel-2, those undergoing post-gelation crosslinking expressed higher levels of *COLX* and *CHAD* (Figure 6B). At the protein level by immunofluorescence (Figure 6C), the expression levels of CILP were similar between constructs with or without interfacial crosslinking, both comparable to the level detected from the native cartilage. In contrast, day 7 stiffening significantly enhanced COLX deposition, with the average MFI value approaching that for the native cartilage. These results indicate that day 7 matrix stiffening promoted the expression of DZ-associated molecules by passaged FTCs.

**Figure 6:**
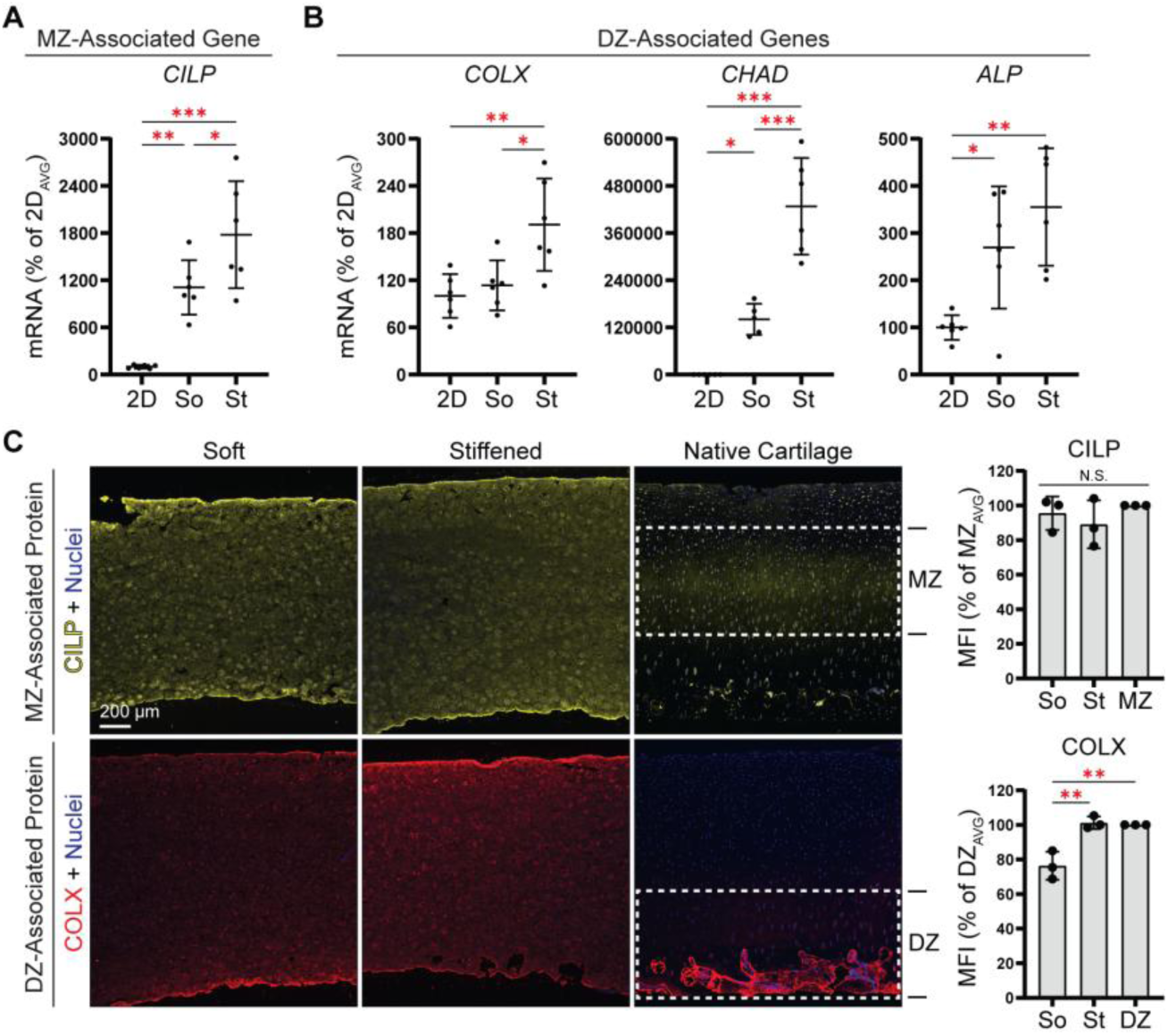
Matrix stiffening during culture increases the expression of DZ-associated proteins by encapsulated FTCs. **(A, B)** Real-time RT-PCR analysis of the expression of MZ- **(A)** and DZ-associated **(B)** proteins by FTCs cultured in soft (So) or stiffened (St) BOHAgel-2. Data are expressed as % of 2D. **(C)** Confocal microscopy images of 3D cultures stained for MZ-associated CILP (Yellow) and DZ-associated COLX (Red). FTCs were maintained in soft (So) and stiffened (St) BOHAgel-2 for 21 days before analysis. Nuclei were counterstained with Hoechst (Blue). The figure on the right shows the corresponding MFI values as a function of culture conditions. Results from native AC were included for comparison. N.S.; Not Significant), *p < 0.05, **p < 0.01, ***p < 0.001.

### Soft hydrogels promote higher expression of PRG4 by passaged SZCs

To determine the effect of hydrogel stiffness on the phenotype of passaged SZCs, we encapsulated monolayer-expanded P2 SZCs in BOHAgel-1 and BOHAgel-2, which were either left unmodified or interfacially crosslinked, thereby generating a range of stiffness conditions. In all conditions, SZCs exhibited predominantly cortical F-actin organization without any detectable protrusions after 21 days of culture (Figure 7A and B). Stiffening BOHAgel-1, but not BOHAgel-2, led to a moderate, but significant, increase in SZC cell area. Cells in both gel types, with or without day 7 stiffening, exhibited similar circularity. At the transcript level, cells in unmodified BOHAgel-1 had the highest *PRG4* expression, although the *CLU* mRNA levels showed no differences across conditions (Figure 7C). To examine whether increases in *PRG4* expression translate into enhanced PRG4 production, we evaluated PRG4 accumulation on day 21 of culture (Figure 7D and E). Again, PRG4 staining and MFI were the highest under the unmodified BOHAgel-1 conditions. Furthermore, PRG4 intensity in soft BOHAgel-1 was similar to that of native SZ cartilage. Thus, the softest condition, BOHAgel-1, promotes native-like SZ expression by passaged SZC.

**Figure 7:**
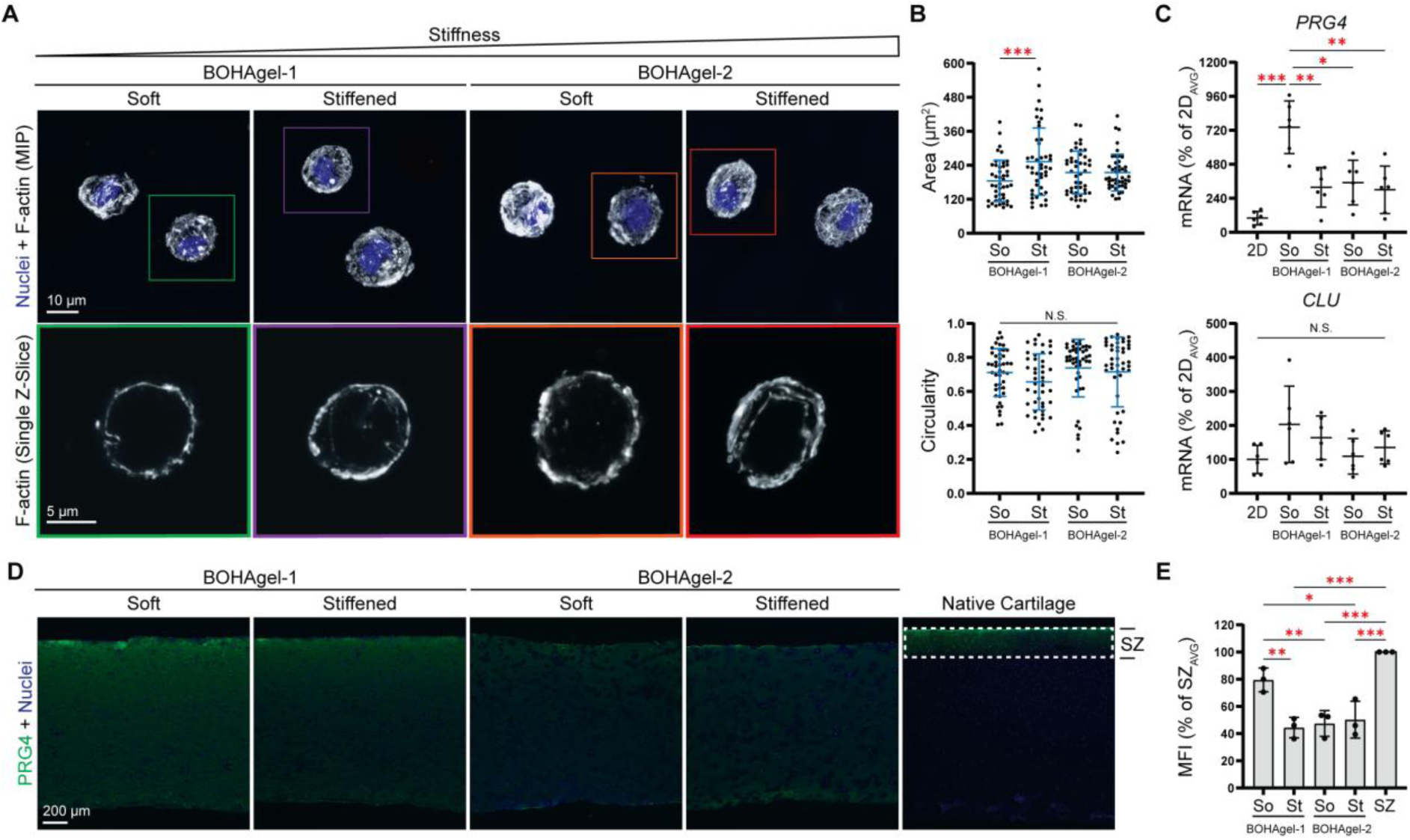
Soft BOHAgel-1 stimulates PRG4 expression by encapsulated P2 SZCs. **(A)** Confocal microscopy images of phalloidin/Hoechst-stained SZCs maintained in BOHAgel-1 or BOHAgel-2 with (Stiffened) or without matrix stiffening (Soft) for a total of 21 days. **(B)** Quantification of cell area and circularity under corresponding conditions. **(C)** Real-time RT-PCR analysis of the transcript levels of SZ-markers expressed as % of 2D controls by encapsulated SZC at day 21. **(D)** Confocal microscopy images of day 21 BOHAgel-1 and −2 constructs with or without stiffening and immunostained for PRG4 (Green). Results from native AC is included for comparison. **(E)** PRG4 mean fluorescent intensity (MFI) as a function of culture condition. N.S.; Not Significant, *p < 0.05, **p < 0.01, ***p < 0.001.

### Multilayered scaffolds with tissue-like stiffness gradient enhance PRG4 and COLX expression in defined SZ and DZ, respectively

So far, we have shown that soft BOHAgel-1 is best suited for SZC culture (high PRG4 expression), whereas stiffened BOHAgel-2 is ideal for FTCs culture. We further demonstrate that dynamic matrix stiffening of BOHAgel-2 promotes upregulated COLX expression. To regenerate the entire chondrocyte population in native articular cartilage with zone-specific phenotypes, we cultured passaged chondrocytes in the multilayered scaffold described above (Figure 2). The construct consisted of a superficial BOHAgel-1 region encapsulating passaged SZCs and the original and stiffened BOHAgel-2 region encapsulating passaged FTCs. These regions comprised approximately 10%, 45%, and 45% of the construct and were designated as the superficial (SZ), middle (MZ), and deep (DZ) zones, respectively (Figure 8A). H&E staining indicates uniform distribution of cells and matrix throughout the construct (Figure 8B). Alcian Blue and Toluidine Blue staining show GAG deposition across all zones, while Picrosirius Red staining depicts collagen deposition throughout the construct, with no apparent zonal differences. Similarly, chondrogenic matrix proteins ACAN and COL2 were expressed throughout all zones, with no significant differences between zones. Importantly, secretion of COL1, a fibroblastic marker, remained low across all zones. However, zonal expression of PRG4 and COLX is obvious (Figure 8D). PRG4 expression is highest in the SZ group compared with the MZ and DZ groups. In contrast, COLX expression is higher in the DZ than in the other zones.

**Figure 8:**
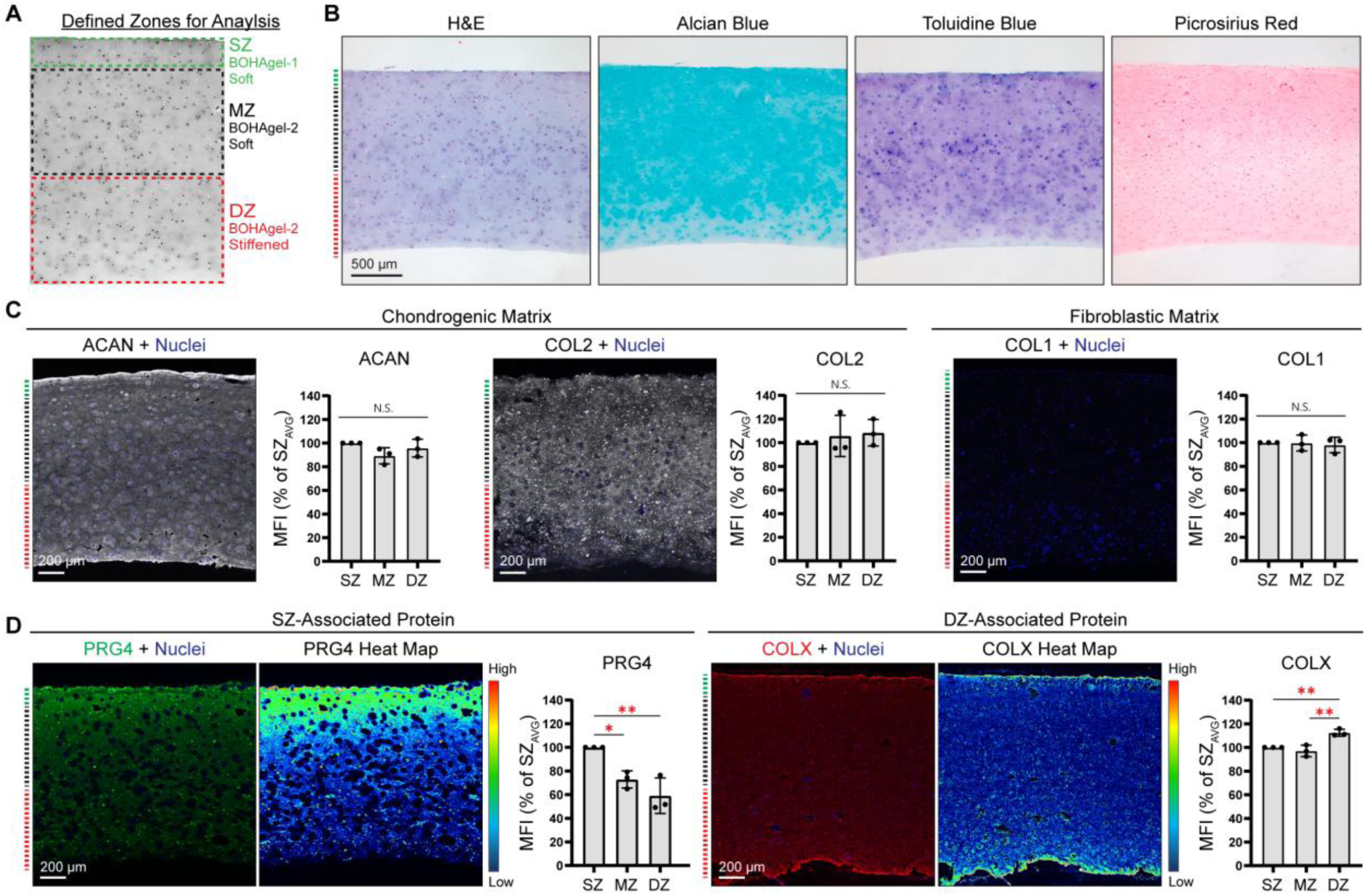
Multilayered scaffolds with zone-specific stiffness enhances expression of PRG4 and COLX in defined zones. **(A)** Representative phase contrast image of the multilayered construct showing defined superficial, middle, and deep zones, corresponding to soft BOHAgel-1, soft BOHAgel-2, and stiffened BOHAgel-2. **(B)** Histological staining of day 21 constructs by H&E, Alcian Blue, Toluidine Blue, and Picrosirius Red. (**C**) Immunofluorescent analysis of the expression of chondrogenic (ACAN, COL2) and fibroblastic (COL1) markers (gray). **(D)** Expression of zone-associated markers PRG4 (Green) and COLX (Red) across the thickness of the gel construct. Samples were counterstained with Hoechst for nuclei (blue). Data are expressed as % of SZ. N.S.; not significant, *p < 0.05, **p < 0.01.

## DISCUSSION

A significant challenge in autologous cell-based therapy for cartilage repair is the differentiation of chondrocytes during *ex vivo* expansion. Moreover, current cartilage repair strategies primarily aim to restore bulk tissue but fail to recapitulate zonal organization and depth-dependent properties, limiting long-term success. Hydrogel-based synthetic matrices with tissue-mimetic zonal architecture offer an attractive strategy for regenerating zonal-specific cell types prior to clinical implementation. The HA-based tunable BOHAgel system directs redifferentiation of passaged chondrocytes throughout culture. Consistent with our previous work on primary mesenchymal and epithelial progenitor cells [34, 36, 46, 47], passaged chondrocytes maintained high viability in custom BOHAgels. Importantly, chondrocyte viability (>88%) exceeds the commonly accepted threshold of 70% [48] for therapeutic applications. Even though the hydrogel displays integrin-binding RGD motifs and is susceptible to MMP cleavage, unlike fibroblasts or mesenchymal stromal cells [34, 36, 46, 47], encapsulated chondrocytes adopted a rounded morphology with cortical F-actin organization, a characteristic feature of native chondrocytes that is lost during monolayer expansion when passaged chondrocytes develop actin stress fibers [49, 50]. Consistent with prior reports on hydrogel-based 3D cultures [6, 25, 51–56], this cortical F-actin organization was accompanied by increased expression of cartilage-specific markers and decreased expression of fibroblastic markers. Upon supplementation with soluble TGFβ3, COL2 expression is rescued, consistent with its reported chondrogenic potential [25, 26, 57]. Collectively, these findings demonstrate that HA-based hydrogels promote redifferentiation of passaged chondrocytes.

While most hydrogel systems used for chondrocyte redifferentiation provide static microenvironments following cell encapsulation, BOHAgels permit user-directed spatiotemporal modulation of matrix stiffness following encapsulation. During cartilage development, a primitive HA-rich soft ECM undergoes rapid mechanical maturation prior to the accumulation of collagen and proteoglycan-rich matrix associated with mature cartilage [31]. The BOHAgel platform used in this study enabled the application of a similar mechanical cue by providing an initially compliant environment followed by *in situ* stiffening. Importantly, the stiffness of these hydrogels remains within a range that supports matrix production, as excessively stiff hydrogel matrices have been shown to act as a barrier to cell activity in 3D [58]. Following monolayer expansion, passaged chondrocytes exhibit a chondrocyte progenitor cell-like (i.e. interzonal) phenotype [27]. Here, we show that matrix stiffening on day 7 of culture increased chondrogenic mRNA expression, promoted rounded cell morphology, and enhanced the production of newly synthesized GAG and collagen. Therefore, providing cells with developmentally inspired cues promotes redifferentiation. This further suggests that stiffness may serve as a cue for regulating zone-specific chondrocyte phenotypes within bioengineered cartilage.

In native cartilage, the SZ, the softest portion of the tissue, is responsible for joint lubrication due to the abundance of PRG4 [6–8], while the DZ, the stiffest layer, contributes to load distribution and integration into the subchondral bone through proteins like COLX [10–12, 15, 16]. We found that hydrogel stiffness played an important role in defining the zonal chondrocyte phenotype. The parent, unmodified BOHAgel-1, the softest gel investigated, promoted an SZ-like phenotype, characterized by increased PRG4 expression by passaged SZC, while stiffened BOHAgel-2, the stiffest gel, enhanced DZ-associated markers, including COLX, by passaged FTCs. The mechanism by which matrix stiffness regulates the zonal chondrocyte phenotype was not directly investigated in this study. Previous studies have shown that matrix elasticity can regulate cell differentiation through mechanosensitive pathways involving integrin signaling, cytoskeletal organization, and mechanically regulated transcription [59], all of which contribute to the maintenance of tissue homeostasis [60]. Chondrocytes primarily sense their local mechanical environment through the pericellular matrix rather than the bulk ECM [61]. The stiffness range examined in this study more closely resembles that of the native pericellular matrix than mature cartilage tissue, supporting the hypothesis that local mechanical environments may regulate zone-specific chondrocyte phenotypes [31].

We further show that spatiotemporal regulation of matrix stiffness may aid in zone-specific cell phenotype and matrix production. We found that combining soft, stiff, and stiffened hydrogel regions within a single scaffold enhanced zone-specific deposition of PRG4 (SZ) and COLX (DZ). To mimic cartilage zonal organization, previous studies exploited the use of gradient maker or relied on gel layering [62–64]. The resulting scaffolds generally exhibit static material properties, contain distinct interfaces between adjacent zones, and do not fully achieve zone-specific protein expression. In contrast, the diffusion-controlled interfacial bioorthogonal crosslinking strategy enabled *in situ* generation of spatially defined microenvironments in a cohesive, integrated cellular construct that more closely recapitulates the depth-dependent organization of native cartilage. With longer culture time and application of mechanical stimulation, this feature can be reinforced by the ECM produced by chondrocyte subtypes residing in each zone

In summary, this study demonstrates that the tunable HA-based hydrogels provide spatially defined microenvironmental cues for 3D culture of passaged chondrocytes. Passaged chondrocytes cultured in BOHAgels and supplemented with TGFβ3 regained the high expression of chondrogenic markers. Matrix stiffening further reinforced the chondrocyte phenotype and stimulated the production of cartilage-specific proteins. We show that hydrogel stiffness played an important role in directing protein expression, with softest hydrogels promoting PRG4 expression and the stiffest one fostering COLX expression. Culturing passaged SZCs and FTCs in an integrated trilayered scaffold with spatially defined stiffness further enhanced expression of zonal proteins in the bioengineered cartilage. Overall, these findings support our hypothesis that tunable HA-based hydrogels can generate zone-specific microenvironments to produce bioengineered cartilage with native-like zonal properties.

Although our work provides a general framework for applying BOHAgel-based strategies to passaged chondrocytes, future studies are necessary to validate this approach using passaged human chondrocytes and determine the translational potential for cartilage repair. While zone-specific marker expression was observed, the engineered tissue likely resembled fetal or newborn cartilage; *in vitro* or *in vivo* tissue development and maturation are necessary to achieve mechanical function. Further studies are needed to evaluate the long-term performance and stability of these constructs to better evaluate their potential for cartilage repair.

## Acknowledgement

This work was supported in part by the Orthopaedic Research and Education Foundation (OREF; 26-010, to AWS, JP, and XJ) and the National Institutes of Health (R01DE029655 and R01DC014461, to XJ). The authors acknowledge the use of facilities and instrumentation supported by the National Science Foundation (NSF) through the University of Delaware Materials Research Science and Engineering Center (DMR 2011824). Microscopy access was supported by grants from the National Science Foundation (NSF; IIA-1301765) and the State of Delaware. This project was supported by the Delaware INBRE program, with a grant from the National Institute of General Medical Sciences (NIGMS; P20 GM103446) from the National Institutes of Health and the State of Delaware. This project was also supported by the DCMR COBRE program, with a grant from the National Institute of General Medical Sciences and from the National Institutes of Health–NIH-NIGMS COBRE (P20 GM139760). We thank Dr. X. Lucas Lu and Annie Porter for their guidance on the metabolic labeling. We acknowledge Genzyme-Sanofi for generously providing HA.

