## Supplementary material for "Bioorthogonal Tuning of Hydrogel Stiffness Promotes Zonal Redifferentiation of Passaged Chondrocytes": Supplemtenal Material

**Synthesis of HA-TCO.**

TCO-hydrazide was prepared by treating TCO-NHS with hydrazine monohydrate, as previously reported [1]. HA (5 kDa, 170 mg, Lifecore Biomedical, Chaska, MN) was dissolved in DI water at 10 mg/mL. To this solution, 1-ethyl-3-(3-dimethylaminopropyl) carbodiimide hydrochloride (EDC, 118.2 mg) and TCO-hydrazide (114 mg, in 1.7 mL of 1/1 mixture of DI water nd DMSO) were added. The reaction was allowed to proceed at room temperature for 3 h, with the pH maintained at 4.75 using 0.1 M HCl. Following the reaction, the pH was adjusted to 8.5 using 0.1 M NaOH, and the mixture was diluted with DI water before being transferred to a dialysis membrane (MWCO: 1 kDa). The product was purified by [2]dialysis against DI water for 3 days, with water exchanged 3 times per day. The purified product was sterile-filtered through a 0.22 µm filter, lyophilized to yield sterile HA-TCO (121 mg, 69.4% yield) as a white, fluffy solid, and stored at -20°C prior to use. The degree of TCO functionalization was confirmed by ^1^H NMR in D_2_O as ~28 mol%.

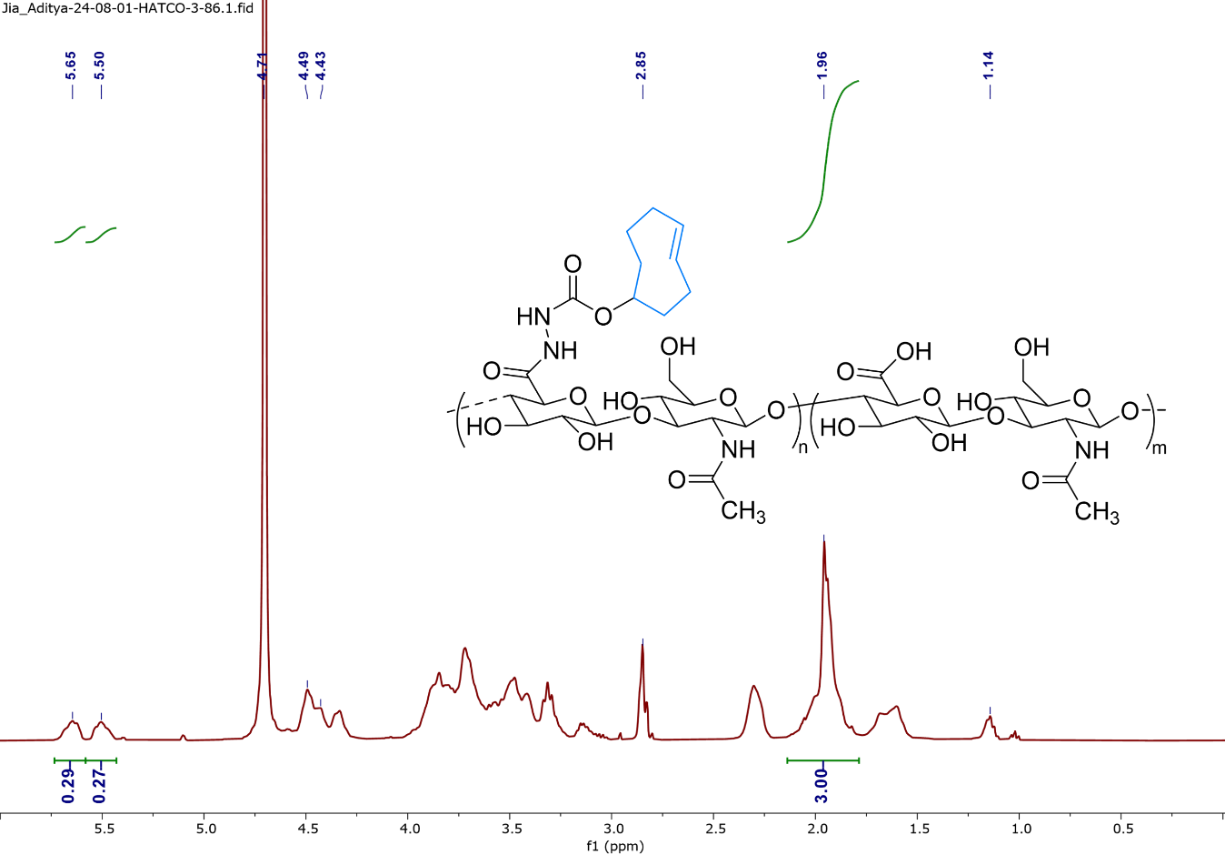

**Figure S1:** ^1^H NMR spectra of HA-TCO in D_2_O.

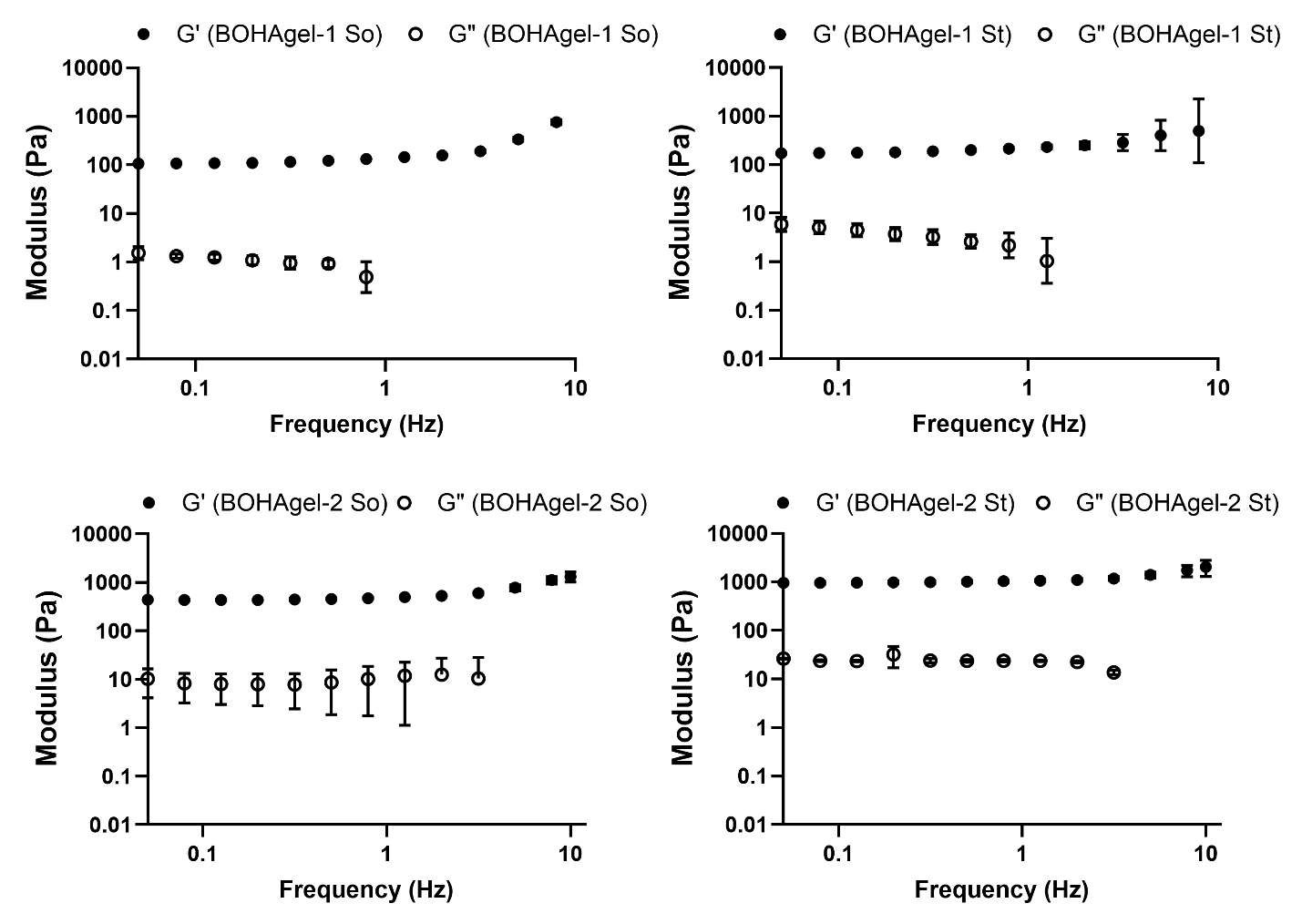

**Figure S2:** Representative frequency sweeps of Orthogel10 and Orthogel20 before (Soft: So) and after stiffening (St).

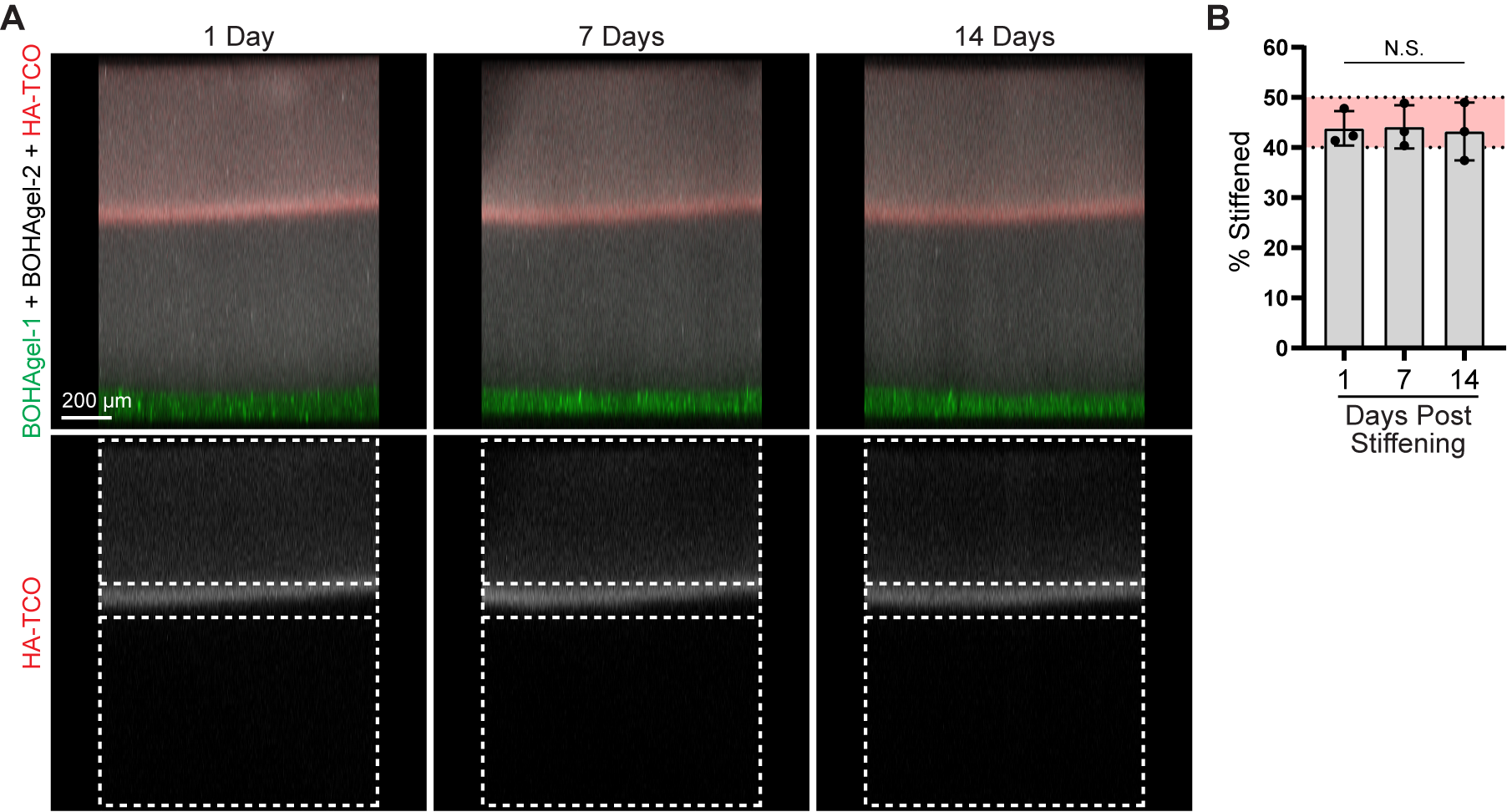

**Figure S3: (A)** Confocal microscopy images of a trilayered hydrogel, consisting of bottom BOHAgel-1 (Green), middle BOHAgel-2 (gray), and top interfacially crosslinked BOHAgel-2 (red), incubated at 37 °C for 1, 7, and 14 days. (**B**) Quantification of the percentage of the hydrogel volume that remained red over time. N.S.; Not Significant.

**Table S1:** Gene-specific primer sequences.

| **Gene** | **Primer Sequence** | **Temp (C)** | **NCBI Reference ID** |
| --- | --- | --- | --- |
| 18S rRNA | **F:** 5’-GTAACCCGTTGAACCCCATT-3’ | 60 | NR_036642.1 |
|  | **R:** 5’-CCATCCAATCGGTAGTAGCG-3’ |  |  |
| Aggrecan  (*ACAN*) | **F:** 5’TGGGACTGAAGTTCTTGGAGA-3’ | 60 | NM_173981.2 |
|  | **R:** 5’-GCGAGTTGTCATGGTCTGAA-3’ |  |  |
| Collagen Type II  (*COL2*) | **F:** 5’-GTGTCAGGGCCAGGATGTC-3’ | 60 | NM_001001135.2 |
|  | **R:** 5’-GCAGAGGACAGTCCCAGTGT-3’ |  |  |
| SRY-Box Transcription Factor 9 (*SOX9*) | **F:** 5’-GTACCCGCACTTGCACAAC-3’ | 60 | NM_001193247.1 |
|  | **R:** 5’-GTGGTCCTTCTTGTGCTGC-3’ |  |  |
| Proteoglycan-4  (*PRG4*) | **F:** 5’-ATGCCTGAACCGACTCCTAC-3’ | 61 | NM_001434744.1 |
|  | **R:** 5’-TGCCGAAGCCTTGACTGG-3’ |  |  |
| Clusterin  (*CLU*) | **F:** 5’-CGGTGACCGAGGCGT-3’ | 60 | NM_173902.2 |
|  | **R:** 5’-TTCCTGGAGCTCTTTGTCCG-3’ |  |  |
| Collagen Type I  (*COLI*) | **F:** 5’-CGGCTCCTGCTCCTCTTAG-3’ | 60 | NM_001034039.1 |
|  | **R:** 5’-CACACGTCTCGGTCATGGTA-3’ |  |  |
| Alpha-smooth Muscle Actin (*αSMA*) | **F:** 5’-ACACCACCCAGAGCGGAGAAGC-3’ | 60 | NM_001034502.1 |
|  | **R:** 5’-GTCCTCCTCCTCGCACATC-3’ |  |  |
| Tenascin-C  (*TNC*) | **F:** 5’-GGAACCTGCGGGCTGTGGAC-3’ | 60 | NM_001078026.1 |
|  | **R:** 5’-CCCCGGATCACCCCGTGGAT-3’ |  |  |
| Chondroadherin  (*CHAD*) | **F:** 5’- AATGCCTTCCAGTCCTTC-3’ | 60 | NM_174019.2 |
|  | **R:** 5’- ACGGTTGTTCTCCAGATG-3’ |  |  |
| Cartilage Intermediate Layer  Protein (*CILP*) | **F:** 5’- TGAGGGGGATCGGTATGACT-3’ | 60 | XM_059890760.1 |
|  | **R:** 5’- GCAAGCCCTGAACTCCATTG-3’ |  |  |
| Collagen Type X  (*COLX*) | **F:** 5’- ACA AGGACCTACAGGAGA AC-3’ | 59 | NM_174634.1 |
|  | **R:** 5’- GCGGCA AGGAGTACA ATG-3’ |  |  |
| Alkaline Phosphatase  (*ALP*) | **F:** 5’- AAGCACTCTCACTATGTCTG-3’ | 60 | NM_176858.2 |
|  | **R:** 5’- GTC GCA TTG TTC CTG TTG-3’ |  |  |

**Table S2:** Immunostaining (A) Primary Antibodies and (B) Secondary Antibodies

| **(A) Primary Antibodies** | | | | | | | |
| --- | --- | --- | --- | --- | --- | --- | --- |
| **Primary Antibody Target** | **Species Isotope** | | | **Dilution** | **Company** | | **Catalog #** |
| ACAN | Mouse Monoclonal IgG1 | | | 1:300 | Thermofisher | | AHP0022 |
| COL1 | Rabbit Polyclonal IgG | | | 1:500 | Abcam | | ab34710 |
| COL2 | Mouse Monoclonal IgG1 | | | 1:300 | EMD Millipore | | MAB8887 |
| COLX | Mouse Monoclonal IgM | | | 1:300 | Abcam | | Ab49945 |
| PRG4 | Mouse Monoclonal IgG1κ | | | 1:300 | EMD Millipore | | MABT401 |
| CILP | Rabbit Polyclonal IgG | | | 1:300 | BIOSS | | bs-13954R |
| **(B) Secondary Antibodies** | | | | | | | |
| **Secondary Antibody Target** | | **Dilution** | **Company** | | | **Catalog #** | |
| CF568 Goat Anti-Rabbit IgG | | 1:500 | Biotium | | | 20103 | |
| CF568 Goat anti-mouse IgG | | 1:500 | Biotium | | | 20100 | |

1.	Gao, H., et al., *Bio-orthogonal tuning of matrix properties during 3D cell culture to induce morphological and phenotypic changes.* Nat Protoc, 2025. **20**(3): p. 727-778.
